# Cell-Cell Separation Device: measurement of intercellular detachment forces

**DOI:** 10.1101/2023.03.16.532950

**Authors:** Julia Eckert, Volha Matylitskaya, Stephan Kasemann, Stefan Partel, Thomas Schmidt

## Abstract

Whether at the intramolecular or cellular scale in organisms, cell-cell adhesion adapt to external mechanical cues arising from the static environment of cells and from dynamic interactions between neighboring cells. Cell-cell adhesions need to resist detachment forces to secure the integrity and internal organization of organisms. In the past, various techniques have been developed to characterize adhesion properties of molecules and cells *in vitro,* and to understand how cells sense and probe their environment. Atomic force microscopy and dual-pipette aspiration, where cells are mainly present in suspension, are common methods for studying detachment forces of cell-cell adhesions. How cell-cell adhesion forces are developed for adherent and environment-adapted cells, however, is less clear. Here, we designed the Cell-Cell Separation Device (CC-SD), a microstructured substrate that measures both intercellular forces and external stresses of cells towards the matrix. The design is based on micropillar arrays originally designed for cell traction-force measurements. We designed PDMS micropillar-blocks, to which cells could adhere and be able to connect to each other across the gap. Controlled stretching of the whole substrate changed the distance between blocks and increased gap size. That allowed us to apply strains to cell-cell contacts, eventually leading to cell-cell adhesion detachment, which was measured by pillar deflections. The CC-SD provided an increase of the gap between the blocks of up to 2.4-fold, which was sufficient to separate substrate-attached cells with fully developed F-actin network. Simultaneously measured pillar deflections allowed us to address cellular response to the intercellular strain applied. The CC-SD thus opens up possibilities for the analysis of intercellular force detachments and sheds light on the robustness of cell-cell adhesions in dynamic processes in tissue development.

## Introduction

Cells are dynamically in contact with each other and with the extracellular matrix (ECM). They experience external mechanical forces applied by neighboring cells and sense, individually or together, changes of the cellular environment, like topography^1^ or stiffness^2^. The passive physical interactions are converted into biochemical signals, which cause an active response followed by an alteration of cell-cell adhesion^3,4^. Cell-cell adhesions have to resist detachment forces^5^. Over the years, various techniques have been developed to characterize the mechanical properties of cells *in vitro* and to understand how cells sense and probe their environment. These techniques, and their further development, for studying cell-cell adhesions have been described in detail in recent reviews^6–13^. In most methodical approaches, like atomic force microscopy^14,15^ and dual-pipette aspiration^16,17^, the detachment forces of cells were studied in a suspended form to avoid cell-ECM interactions. In the experiments, cells were brought into contact for a short period of time such that they did not develop any actin stress fiber network. Using other techniques such as Förster resonance energy transfer (FRET)-based tension sensors, classical traction force microscopy (TFM), and micropillar-array traction technology cells are investigated that adhered to substrates. Those experiments give an insight into the force and tension interactions of cells ^18–23^. The intercellular forces are predicted to be dependent on the ECM-dependent traction forces, on cell spreading area and shape, on cell-cell contact size, and on the cell type with its different cell-cell adhesion machinery.

Here, we combined ideas of measuring intercellular detachment-forces of cells in the spread state that have developed the shape-determining actin stress fibers in a single device. The Cell-Cell Separation Device (CC-SD) was able to obtain intercellular forces through traction force measurements of deflected micropillars. In addition, the micropillars were constructed on top of thicker PDMS-blocks that were separated by few micrometer-sized trenches or gaps. Cells adhered to the top of the micropillars and were able to connect across the gap. This device was mounted onto a linear stretcher device by which a controlled stretch on a thinner homogeneous substrate underneath the blocks was applied, and the distance between the cells was increased until a cell-cell contact broke. We here demonstrate that the CC-SD provided a nominal strain at the cell-cell contact by an increase of the gap width between the blocks by up to 140% (i.e. 2.4-fold) which was sufficient to separate substrate-attached cells with fully developed actin stress fiber network. Simultaneously, pillar deflections caused by cell traction forces gave information about the cellular responses to the intercellular strain. The design opens up possibilities for the analysis of intercellular force detachments, and sheds light on the robustness of cell-cell adhesions in dynamic processes that are at the basis of tissue development.

## Results

### The intercellular adhesion strength increases with an increase of the total traction force

Cells form doublets (Fig.1A), larger cell clusters up to confluent monolayers, ultimately forming tissues that define the organism. The stability of such cellular assembly is secured by the interplay of cell-matrix and cell-cell adhesions ^4^. At the single-cell level, where solely cell-matrix adhesion is important, it has been shown that the summed absolute traction force that individual 3T3 fibroblast cells exert on the extracellular matrix scales with the cell spreading area. Accordingly, for cells growing on fibronectin-coated micropillars, the total absolute traction forces increased proportional to the number of deflected pillars below the cell ^24,25^. The question we asked here was, whether this linear scaling between interaction area and total absolute force prevails for cell assemblies, here in particular for cell-doublets. In order to test the hypothesis, we cultured cells on flexible micropillars and analyzed the traction forces developed by cells after fixation. Experiments were performed on 46 cell-doublets (Fig.1B-C). The total absolute traction force linearly increased with the number of deflected pillars for both the cell doublets and the 92 individual cells after forces were split and assigned to each cell separately. This relationship was identical for the doublet configuration as well as for individual cells (Fig.1D). The linear dependency suggest that the mean absolute force per deflected pillar is a constant that reflects a property of the individual cell. The mean absolute force per deflected pillar was calculated to be 11.0 ± 2.7 nN for doublets, and 10.9 ± 3.4 nN (mean ± s.d.) for individual cells of 3T3 fibroblasts on fibronectin-coated pillars. Similar results have previously been reported for individual 3T3 fibroblasts^25^.

**Fig. 1.**
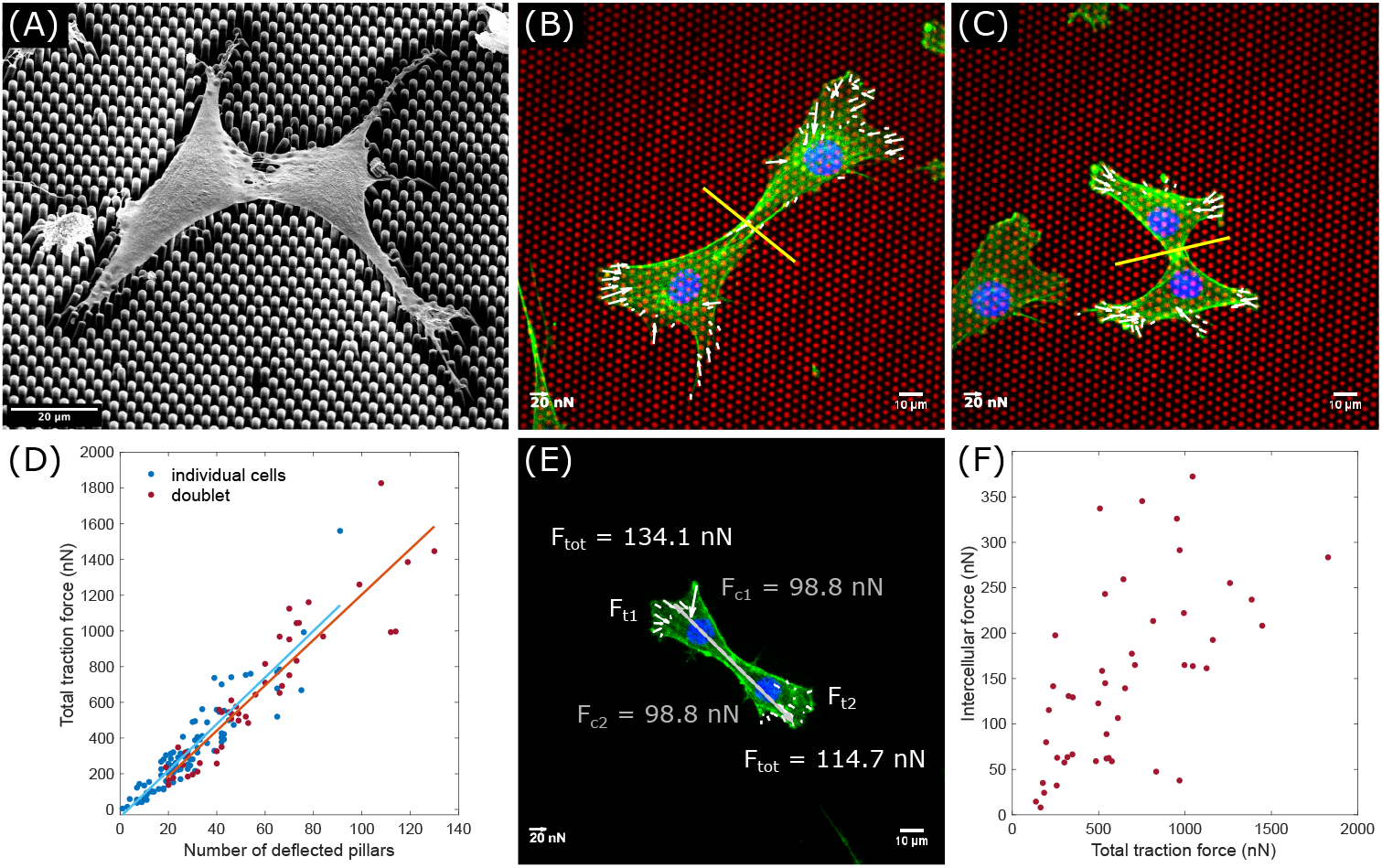
The intercellular force increases with the total traction force. (**A**), Scanning electron microscopy image of a 3T3 fibroblast cell-doublet. (**B,C**), Two 3T3 cells connected to a cell-doublet exert traction forces (white arrows) on elastic micropillars. For the traction force analysis of each individual cell, the doublet was separated at the cell-cell contact (yellow line). Red: fibronectin-coated micropillars, green: F-actin, blue: nuclei. (**D**), Total traction force, Ftot, per doublet and for each individual cell versus the number of deflected pillars. (**E**), Traction forces, *F_t_* of each individual cell add up and result in the intercellular force contribution, *F_c_*. (**F**), The intercellular force, *F*_cc_ = |*F*_c1_| + |*F*_c2_|, versus the total traction force per doublet. Correlation coefficient: r = 0.6.

The above measurements on cell-doublets allowed us to measure the forces that developed at the cell-cell contacts^19^. After partitioning the cell-doublet into two individual cells, we calculated the resultant force, *F*_c_, for each cell, c1 and c2, by summing up the individual traction forces, *F*_t_ (Fig.1E; Eqs.(7)). The resulting forces *F*_c1_ and *F*_c2_ were opposing, as required from Newton’s law. Our results showed that the intercellular force (Eq.(8)) increased with the magnitude of the total traction force exerted by the doublet (Fig.1F). About 25% of the total traction force was accounted for the force between cells. Therefore, we speculate that 25% of the active adhesion molecules are located at the cell-cell interface.

### Stretching a homogeneous elastic micropillar field causes deformations

In an organism, the static picture we analyzed in the previous subsection is falling short. The forces between cells, and between cells and the extracellular matrix constantly change due to the cell’s activity. Hence, cell-cell adhesion needs to adapt in a dynamic way to keep tissue integrity intact. To test how cell-doublets and their individual cells adapt to dynamic mechanical challenges, we designed an experiment that included a dynamic stretch of the total elastic micropillar array. Earlier, it has been shown that the cell’s cytoskeleton largely rearranged on a continued application to uniaxial stretch^26^. Here, we used a similar approach to assess whether a short uniaxial stretch would lead to a change in cell-cell adhesion forces, and whether we would be able to determine the forces that are sufficient to break a cell-cell contact. Our experimental protocol was performed in less than two minutes to avoid intracellular responses such as cytoskeleton rearrangements.

We cultured human melanoma MV3 cells on micropillar arrays and mounted arrays of thickness of of ~100 μm with two clamps in a linear stretcher on a confocal microscope. The clamp-positions were electronically controlled (Fig.2A). We first characterize the resulting strain-fields that develop on homogeneous pillar arrays by analyzing the pillar positions with respect to the external strain applied. At 0% strain, i.e. in an unstretched position, the mean pillar-to-pillar distances was 3.87 ± 0.18 μm (mean ± s.d.) as predicted from the silicon master (Fig.2B,D). When we applied a stretch in x-direction on the micropillar array, we identified a displacement of the pillars in both the x- and y-directions (Fig.2B-C). At 20% nominal strain, the pillar-to-pillar distance increased to 4.05 ± 0.15 μm, i.e. 4.6% in the stretch direction, and decreased to 3.79 ± 0.09 μm (mean ± s.d.), i.e. −2.3% perpendicular to it (Fig.2D). The x-y difference is predicted for an incompressible material like PDMS, for which the Poisson ratio *v* = 0.5, and for which the strain relation (Eq.(6)) holds.

**Fig. 2.**
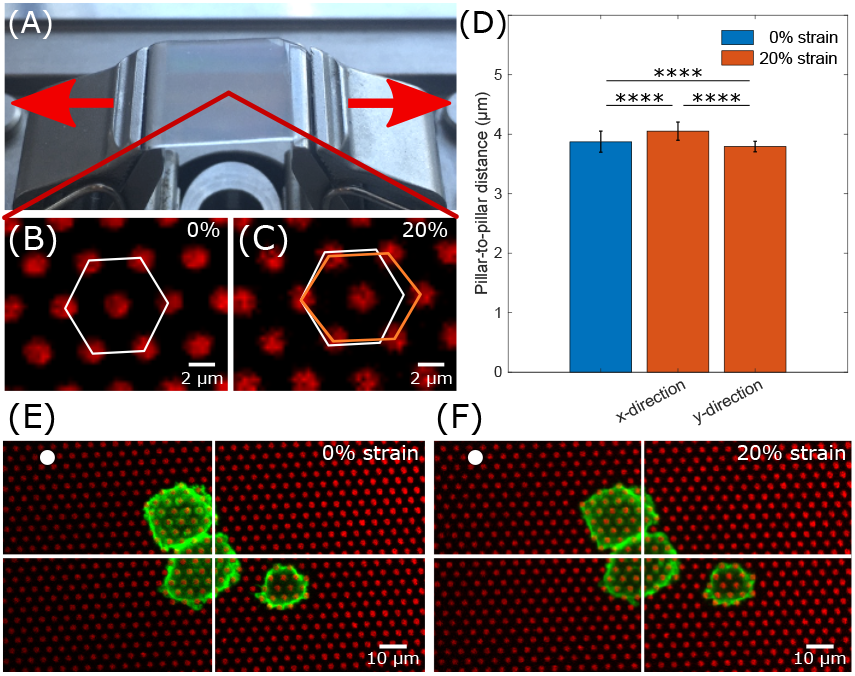
x-y displacement changes by less than 5%. (**A**), A 1×1 cm micropillar array mounted with two clamps. The horizontal stretch direction of the array is indicated by red arrows. (**B**), Top view of a micropillar with hexagonal geometry in the unstretched position. (**C**), A strain of 20% on the pillar array caused a deformation of the substrate and thus an increase and decrease of the center-to-center distance in the x- and y-position, respectively, which is indicated by the orange hexagon. Scale bar: 2 μm. (**D**), Pillar-to-pillar distances at 0% and 20% substrate strain in x-direction (n=40). (**E,F**), MV3 cells adhered to micropillars undergo substrate stretching and maintain stable cell-cell adhesions. The white dot represents the reference point of both images. Red: fibronectin-coated micropillars, green: F-actin. Two-sided Wilcoxon rank sum test: **** p < 0.0001.

It is essential for the methodology of force measurements in micropillar assays that the positions of the undeflected pillars are known to high accuracy ^27^. When stretching the substrate, this accuracy might be deteriorated by, for example, pillar deformations and imperfections in the homogeneity of the PDMS polymer. Thus, we further characterized the background deflection field, i.e. excluding pillars covered by cells, of the micropillars on uniaxial stretch. For an unstretched array, the mean deflection was 0.038 ± 0.021 μm (mean ± s.d.) (Fig.S2A). This accuracy was predicted from the total signal detected for each pillar on the CCD-detector of the microscope^27^. At 20 % nominal substrate strain, the mean background deflection increased by 38% to 0.061 ± 0.033 μm (mean ± s.d.) (Fig.S2B-C).

To check whether the applied substrate strain affected the traction force of cells, we focused on the pillar deflection analysis of single MV3 cell. These control experiments are needed in order to rule out the mechanical influence of substrate deformation on the intercellular force calculation. At 0% nominal strain, the mean deflection of the pillars was 0.32 ± 0.17 μm (mean ± s.d.) and thus an order of magnitude higher than the background deflections (Fig.S2D). When the 20% nominal strain was applied, the mean deflection of 0.34 ± 0.26 μm (mean ± s.d.) did not change compared to the unstretched position (Fig.S2E-F). The stretch of the micropillar and thus the deformation of the pillar field underneath the cell did not affect the mean traction forces of the MV3 cell.

These experiments demonstrated that the micropillar technology can be used to apply a defined stress/strain on both cells and cell assemblies, with the ability to simultaneously monitor the cellular force-response as monitored by the micropillar deflections. Yet, likewise, the results showed a severe limitation of this initial approach: the local strain field, reflected by the pillar-to-pillar displacement, that could be achieved was less than 5% on a nominal substrate stretch of 20%. Probably most of the strain was localized to areas of the substrate weakened by the clamps of the stretcher. The amount of strain that was realized on cells was not sufficient to challenge a cellular response, or even break the cell-cell contacts as shown for the MV3 cells in Fig.2E-F.

### CC-SD: the Cell-Cell Separation Device

In order to break the cell-cell contact and simultaneously measure the maximum intercellular adhesion force between two cells adhered to the substrate when in a doublet configuration, we designed a novel substrate, the Cell-Cell Separation Device (CC-SD) (Fig.3A). The substrate is composed of PDMS blocks of height, *H*, connected by a thin layer of PDMS at the bottom. On top of each block, a field of micropillars is located to which cells can adhere. The blocks are spaced in such a way that cells are allowed to connect across the gap of width, *R* (Fig.3E). An externally applied stretch on the thin layer localizes the strain to the gap and increases significantly the block distance, *R*. This eventually breaks the cell-cell contact.

**Fig. 3.**
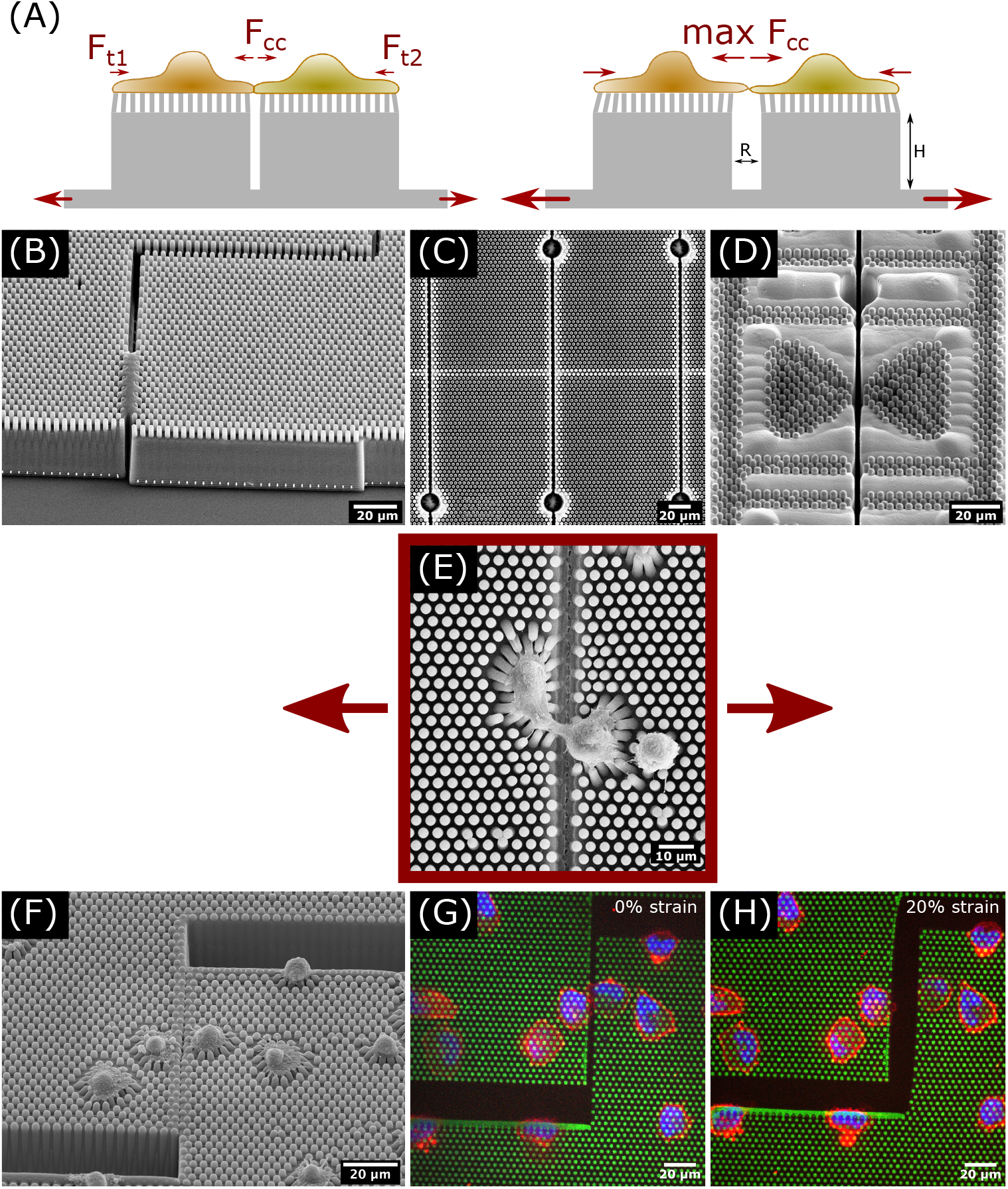
Cell-Cell Separation Device (CC-SD). (**A**), Schematic overview of the CC-SD. Two cells adhered on blocks of height, H, composed of micropillars and connect across a gap of width, R. Obtained traction forces, *F_t_*, (Eqs.(7)) result in the intercellular forces, *F*_cc_ (Eq.(8)). By applying a stretch on the connected substrate, the gap width increases, resulting in an increase in pillar deflections, i.e. increase in pseudo-traction forces. At a certain strain, the cell-cell adhesions break and the maximum intercellular force can be obtained. The size of the red arrows indicates the strength of the force. (**B**), CC-SD in a zig-zag configuration for tensile and shear force measurements at the cell-cell contact. (**C**), Parallel gaps perpendicular to the stretch direction. (**D**), CC-SD with pillar fields in a butterfly configuration. The geometry of the wings controls the cell spreading area and contact length between the cells. (**E**), Two MV3 cells adhere across the gap. (**F-H**), CC-SD in a zig-zag configuration. An application of 20% strain is sufficient enough to break the cell-cell contact of MV3 cells. Green: fibronectin-coated micropillars, red: F-actin, blue: nuclei.

Our novel design, although adding substantial challenges in the microfabrication process by deep reactive-ion etching (see materials and methods), allowed us to develop additional application modes, which make the CC-SD a versatile instrument. We developed and characterized four CC-SDs in different configurations (Fig.3B-D,F) whose specifications are summarized in Tab.S1. In one design, the gap was arranged perpendicular to the direction of stretching. That design allowed to apply tensile stress at the cell-cell contact during uniaxial stretching (Fig.3C). In a second and third design, the gap was arranged in a zig-zag pattern, allowing additional measurements of shear forces where the blocks separated parallel to the cell-cell contact length (Fig.3B). The pillars in the fourth design, inspired by Liu et.al.^19^, were arranged in a butterfly shape to adjust the cell spreading area and allowed the cell-cell contact length to be fixed (Fig.3D).

All CC-SDs were designed to be optimized for the cell lines we use in ongoing studies: the 3T3 fibroblast cell line and the MV3 melanoma cell line. As in the design of the homogeneous arrays we used earlier, the micropillar fields on top of the blocks consisted of pillars of ~2 μm diameter arranged in regular hexagonal patterns of ~4 μm center-to-center distance. The pillar height was adjusted to the range of suitable effective Young’s moduli relevant to the two cell types. For a pillar height of 6.1 μm of the previously used micropillar array (effective Young’s modulus: 29.5 kPa), fixed 3T3 fibroblasts were reported to apply a mean force per pillar of 13 ± 5 nN (mean ± s.d.)^25^. We further measured a mean traction force of life 3T3 fibroblasts of 15 ± 7 nN, which is 13% more than in the fixed condition, and life MV3 cells of 19 ± 6 nN (Fig.S1A). Using pillar heights of 4.1 μm (effective Young’s modulus: 47.2 kPa), the mean traction forces of MV3 cells reduced to 11.0 ± 3.5 nN (Fig.S1B). The initial gap width of all CC-SD substrates was set to 4 μm perpendicular to the direction of stretching, a length scale across which 3T3 and MV3 cells were able to span and adhere to each other (Fig.3E; Fig.4A). The area of the butterfly pattern was chosen to match the mean spreading area of the MV3 cell line, which was between 117 ± 61 pillars and 122 ± 85 pillars (mean ± s.d.) (Fig.S1C). Accordingly, we chose an area of each butterfly wing of 119 pillars.

**Fig. 4.**
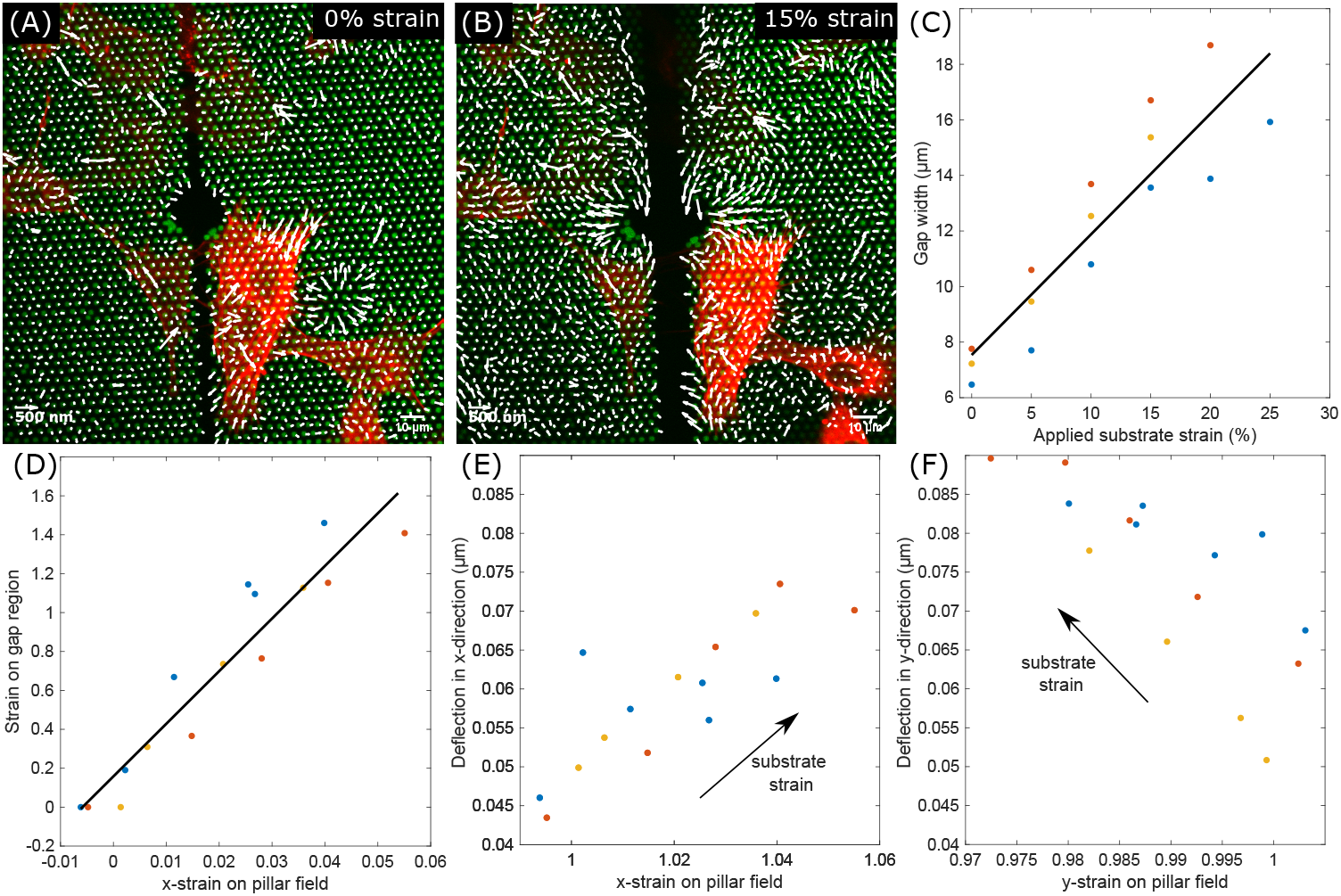
Tensile CC-SD. (**A,B**), CC-SD with the gap perpendicular to the stretch direction, showing 3T3 fibroblasts and the pillar deflection field (white arrows) at 0% and 15% substrate strain. Green: fibronectin-coated micropillars, red: Factin. (**C**), By applying a strain on the substrate, the width of the gap increases (n = 3). (**D**), The strain on the gap region is 27.04 ± 2.54 bigger than the deformation on the pillar field. (**E,F**), The magnitude of the computed pillar deflections in x- and y-directions increase with increasing substrate strain. The stain on the substrate causes an elongation (increased strain) of the pillar field in x-direction (**E**) and a compression (decreased strain) in y-direction (**F**).

We set the block height between 30 μm to 40 μm (Fig.3B). This height was sufficient to avoid pre-stretching our substrate while mounting. The strain on the CC-SD was mainly localized and applied in the region between the blocks. When we stretched the tensile CC-SD with the perpendicular gap, we identified a linear increase of the gap width of 0.43 ± 0.06 μm per nominal strain, i.e. from ~6 μm at 0% stretch to ~16 μm at 20% nominal stretch (Fig.4C). Furthermore, we compared the strain in x-direction of the pillar field with the strain on the gap region (Fig.4D). The gap width increased by 27.04 ± 2.54 times the pillar-to-pillar distance. In comparison to the tensile stretch of 4.6% of the pillar array (Fig.2D-F), the gap width increased by 140.2%, i.e. 2.4-fold.

As shown for the micropillar array above, the substrate deformation caused by the nominal substrate strain affected the accuracy of pillar detections. We showed that the magnitude of the pillar deflections in x- and y-direction increased with increasing substrate strain (Fig.S2C). We wanted to confirm whether the earlier result likewise holds for the CC-SD substrate. For that, we computed the pillar deflections for each nominal substrate strain applied (Fig.4E-F). We found an increase of the pillar deflections in x- and y-direction with increasing nominal stretch, which further correlated with the elongation and compression of the substrate, respectively. One example of 3T3 fibroblasts cultured on the tensile CC-SD is shown in Fig.5A-B. Two cells adhered to each other across the gap and were stretched by increasing gap width. The mean deflections of the background were 0.0481 ± 0.0022 μm and 0.0965 ± 0.0039 μm (mean ± s.e.m.) at 0% and 15% nominal strain, respectively (Fig.5C). When the stress at the cell-cell contact increased, we expected a cellular response by increasing pillar deflections towards the gap to obey Newton’s third law in terms of total force conservation. In order to check whether the strain at the cell-cell contact affected the overall applied traction forces, we analyzed the pillar deflections below the cells (Fig.5D-E). At 0% strain, the mean deflection caused by the cells was found to be 0.070 ± 0.006 μm (mean ± s.e.m.). When we applied the nominal substrate strain of 15%, the mean pillar deflections increased to 0.137 ± 0.009 μm (mean ± s.e.m.) (Fig.5F). Since the mean pillar deflections, 〈*δ*_all_〉, below the cells were close to the mean background deflections, 〈*δ*_bkg_〉, we needed to correct for the background. Assuming that the real cellular deflections, 〈*δ*_cell_〉, are uncorrelated to the background as given by the positional accuracy, the real mean deflection is given by 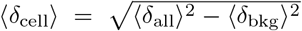 (Eq.(4), Eq.(5)). This value changed from 〈*δ*_cell_〉 = 0.050 ± 0.008 μm at 0% to 〈*δ*_cell_〉 = 0.098 ± 0.013 μm at 15% substrate strain, indicating an increased strain at the cell-cell contact during stretching according to Eqs.(7). A similar result is shown for a single MV3 cells in Fig.S3. We measured an increase in pillar deflections in the x-direction when we applied the stretch to the substrate. Unfortunately, the pillar height decreased towards the gap due to technical limitations in the manufacturing of the Si-wafer for the PDMS substrate. This challenge needs to be tackled to allow us to translate deflections into forces in the future.

**Fig. 5.**
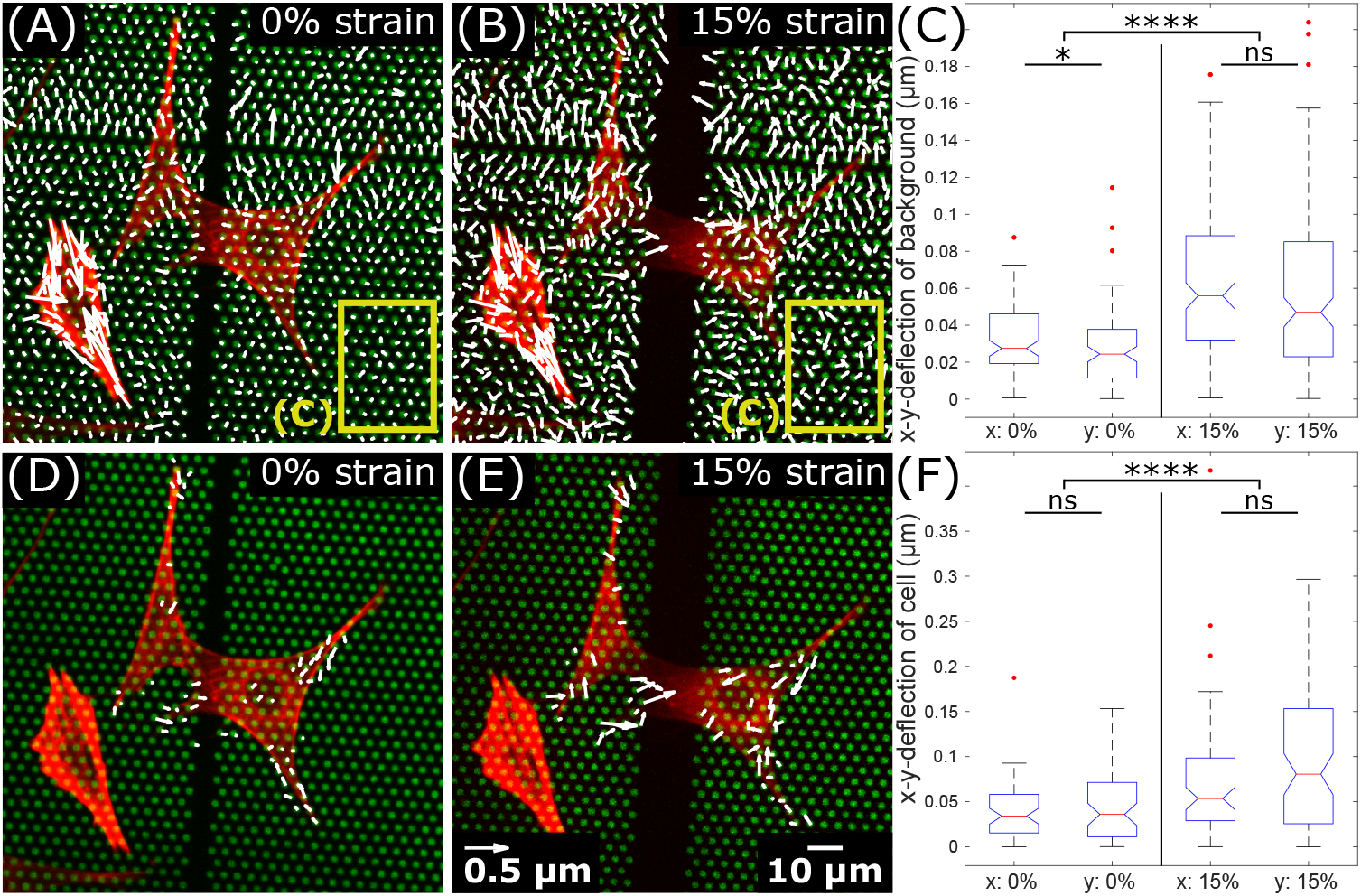
Traction forces of cell doublet increases with increasing gap width. 3T3 fibroblast cells cultured on the CC-SD with the gap perpendicular to the direction of stretching. (**A,B**), The deflection field of all pillars at 0% and 15% nominal substrate strain. The yellow frame represents the pillars considered for background analysis in (**C**). (**C**), Absolute deflections in x-and y-direction of pillars in a manual selected background region, i.e. excluding pillars covered by cells. In total, 98 and 154 pillars at 0% and 15% nominal strain, respectively, were considered. (**D,E**), Deflection of pillars caused by cell traction forces at 0% and 15% nominal substrate strain. All deflections pointing towards the center of the cell are shown and were used for the analysis. (**F**), Absolute deflections in x-and y-direction of pillars deflected by the cell. In total, 58 and 77 pillars at 0% and 15% nominal strain, respectively, were considered. Green: fibronectin-coated pillars, red: F-actin. Two-sided Wilcoxon rank sum test: * p ≤ 0.05, **** p < 0.0001, ^*ns*^ p > 0.05.

## Discussion

Our goal in the present study was to provide a design for intercellular force detachment measurements of substrate-attached and spread cells that were allowed to fully develop an actin stress-fiber network prior to experimentation. The design of a structured force-sensor field does allow us now to measure, in real-time, the cellular force response on an applied linear stress at the cell-cell contact. The strain applied can be up to 140%, i.e. 2.4-fold. This large increase in strain was sufficient to successfully break cell-cell adhesions of MV3 cells (Fig.3G-H) and of 3T3 fibroblasts (Fig.4A-B). By localizing the stress to the cell-cell contact, cells themselves were unaffected by substrate deformations, unlike in earlier cell-stretching methodologies^28,29^.

In using the CC-SD to pull cells apart from each other, stress will be built up at the cellcell contact. Simultaneously, for each single cell and cell doublet, the law of conservation of forces holds. Thus, we expected a change in the pillar deflection field below the cells as long as they were in contact (Fig.5F). The increased tension at the cell-cell contact thus is predicted to increase the average magnitude of the total pillar deflections per cell doublet. Further, we predicted a constant number of deflected pillars for all applied substrate strains. These predictions at least hold for shortterm mechanical challenge. It has been previously reported that force-exertion by cells follows intracellular signalling process by a time delay of about three minutes ^30^. Our experiments were performed at shorter timescales of about two minutes.

It has been reported earlier that traction forces of cells act parallel to the cell’s elongation ^25^. Indeed, our data on the elongation of both 3T3 fibroblasts showed traction forces developing parallel to the gap, i.e. perpendicular to the direction of substrate stretching. Thus, our findings corroborate such predictions. Our novel design opens up possibilities for the analysis of intercellular force detachments in the future, and sheds light on the robustness of cell-cell adhesions, eventually during the dynamic processes in the development of tissues.

## Methods

### Cell culture

3T3 fibroblast and MV3 cells were cultured in high-glucose Dulbecco Modified Eagle’s Medium (D6546; Sigma-Aldrich, St. Louis, MO, USA) supplemented with 10% fetal calf serum (Thermo Fisher Scientific, Waltham, MA, USA), 2 mM glutamine, and 100 mg/mL penicillin/streptomycin, 37 °C, 5% CO_2_.

### Immunostaining

For experiments on homogeneous pillar arrays, after 22.5 h of spreading, 3T3 fibroblast cells were fixed for 15 min in 4% paraformaldehyde (43368; Alfa Aesar, Haverhill, MA, USA) in phosphate-buffered saline (PBS). Furthermore, cells were permeabilized for 10 min with 0.1% Triton-X 100 in 1% bovine serum albumin (BSA) and blocked for 60 min with 1% BSA in PBS. F-actin was stained with Alexa Fluor 532-labeled phalloidin (A22282; Invitrogen, Carlsbad, CA, USA) and the DNA with DAPI (Sigma-Aldrich).

### Life-cell staining

2 h before imaging, Factin of MV3 cells was stained with CellMask dye (A57245; Invitrogen, DeepRed Actin) and DNA with Hoechst 34580 (63493; Sigma-Aldrich).

### Wafer manufacture of CC-SD substrate

The CC-SD substrate was manufactured in several dry etching steps (Fig. S4). In a first step, a silicon-wafer of 100 mm in diameter was oxidized by thermal oxidation (SiO_2_ thickness: 430 nm). Afterwards, the first photolithography step took place, in which a thin layer of photoresist (PR; MEGAPOSIT SPR 955CM 0.7) was utilized to transfer the desired pattern, and open the SiO_2_ mask. In this step, the holes that will become the PDMS pillars later were defined. After opening the SiO_2_ hard mask, the holes were transferred into the Si-substrate using the Bosch process to etch deep into the substrate. The remaining photoresist was removed and another SiO_2_ hard mask created by thermal oxidation. Next, the second photoresist layer was applied by spraying. This layer was used to separate the hole arrays (block height, *H*) followed by opening the oxide mask by deep etching using the Bosch process. Precise coordination of the etching depth of the two dry etching steps was essential, since this defines the height of the pillars. This depth is strongly related to the diameter of the holes. In the final step, the remaining silicon oxide was removed by wet etching (BOE).

### Elastic micropillar arrays and CC-SD

Polydimethylsiloxane (PDMS, Sylgard 184) micropillar arrays of 2 μm diameter, 6.1 μm (E_eff_ = 29.5 kPa) and 4.1 μm length (E_eff_ = 47.2 kPa), and 4 μm spacing in a hexagonal geometry were used for cell traction force experiments. The pillar arrays were flanked by 50 μm spacers on two sides of the array. Details of this arrangement and the experimental procedures were described earlier in detail^27^. In brief, pillar arrays were produced on a negative silicon-wafer master made by a two-step deep reactive-ion etching process. Wafers were passivated in trichloro-silane (448931; Sigma-Aldrich). A mixture of 1:10 PDMS (cross-linker/base ratio) was poured onto the Si-master and cured for 20 h at 110 °C. Thin CC-CD substrates of about 100 μm were peeled off in absolute ethanol for subsequent critical point drying. The tops of the pillars were coated by micro-contact printing. For that, flat 1:30 PDMS stamps were incubated for 1 h with 40 mL of 50 mg/mL Alexa Fluor 647–labeled or Alexa Fluor 532-labeled, and 50 mg/mL unlabeled fibronectin (F1141; Sigma-Aldrich), then washed and dried. Subsequently, the stamps were gently loaded onto the ultraviolet-ozone-activated micropillar arrays for 10 min. After stamping, the arrays were passivated with 0.2% Pluronic (F-127, P2443; Sigma-Aldrich) for 1 h, and washed in PBS.

### Stretcher

We used an in-house made, piezo-driven stretcher ^26^. PDMS samples of 100 μm thickness were mounted with two clamps (Fig.2A). A uniaxial stretch to the PDMS layer was applied by two independent piezo motors (SLC2430s, SmarAct) and a controller unit (MCS-3D, SmarAct, Oldenburg, Germany). An in-house written LabVIEW program was used to control the strain and strain rate. For our experiments, we took images ever 5% strain and increased the strain by a rate of 0.5 %/s.

### Microscopy

Samples were imaged at high resolution on a home-build optical microscope setup based on an inverted Axiovert200 microscope body (Carl Zeiss, Oberkochen, Germany), a spinning disk unit (CSU-X1; Yokogawa Electric, Musashino, Tokyo, Japan), and an emCCD camera (iXon 897; Andor Labs, Morrisville, NC, USA). IQ-software (Andor Labs) was used for setup-control and data acquisition. Illumination was performed using fiber-coupling of different lasers (405 nm (CrystaLaser, Reno, NV, USA), 514 nm (Cobolt AB, Solna, Sweden), and 642 nm (Spectra-Physics Excelsior; Spectra-Physics, Stahnsdorf, Germany)). 3T3 cells on pillar arrays for doublet analysis were placed upside-down onto 25 mm cover glasses and viewed with an EC Plan-NEOFLUAR 40 × 1.3 Oil Immersion Objective (Carl Zeiss). The stretcher unit for cell-cell separations was mounted on the microscope.

### Scanning electron microscope

After cell fixation and washing steps with PBS buffer, the medium was gradually replaced by a mixture of ethanol and milliQ water: 50%, 70%, 80%, 90%, and 100%. Each incubation step was 10 min. Next, the sample was transferred in 100% ethanol into the chamber of the critical point dryer (CPD 020, Balzers). After the replacement of ethanol with liquid CO_2_, the samples were dried. Before the investigation with the scanning electron microscope, samples were coated with lead/platinum.

### Pillar stiffness characterization

The relation between traction force to pillar deflection was calculated from Euler’s beam theory for flexible beams, supplemented by correction terms^31^. The total pillar deflection was described by bending, shear, and tilting deflection as

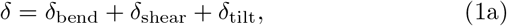

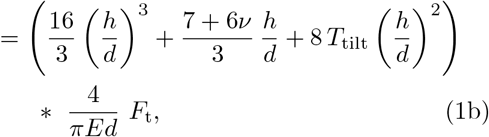

where *F*_t_ is the traction force, *E* is the Young’s modulus, *d* is the pillar diameter, *h* is the pillar length, *v* is the Poisson ratio, and

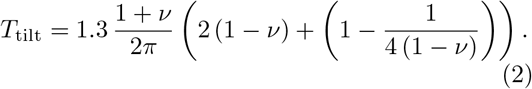

is the tilting coefficient^31^. For our experiment, *v* = 0.5 and E = 2,500 kPa for PDMS. Accordingly, the traction force applied on the pillar was *F_t_* = *kδ.*

The effective Young’s modulus of the pillar field, in relation to that of a homogeneous object, is given by^32^

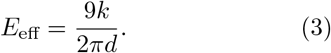

### Image analysis

Images of MV3 and 3T3 cells within the field of view of 176 × 176 μm were analyzed using MATLAB scripts (MATLAB R2018a; MathWorks, Natick, MA, USA). Pillar deflections were quantified as previously described in detail^27^. Deflected pillars, caused by cell traction forces, were distinguished from the background deflections by thresholding: the distribution of background deflections were determined from an undeflected area of the pillar array by selecting a pillar region outside the cell area. Pillar deflections underneath the cell that fall within the so-constructed background range were excluded.

For cells in the CC-SD images, the mean of cellular deflections was calculated by

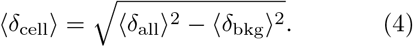

Here, 〈*δ*_all_〉 is the mean of all deflections below the cell and 〈*δ*_bkg_〉 is the mean deflection of the background. The error of 〈*δ*_cell_〉 was derived from error propagation

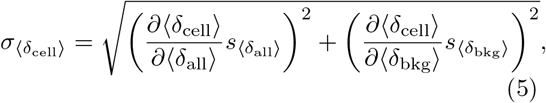

with *s*_〈*δ*_all_〉)_ and *s*_〈*δ*_bkg_〉_ the standard error of means.

### Poisson correction

To determine the pillar deflections, we considered the x- and y-deformation of the substrate caused by the uniaxial stretch by using the Poisson ratio:

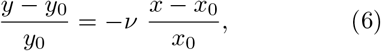

with *x*_0_, *y*_0_ the unstretched and *x, y* the stretched positions.

### Intercellular force analysis

The intercellular force contribution of each cell in a doublet configuration was calculated by

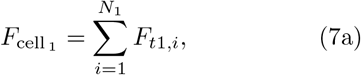

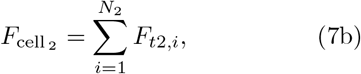

where *F_t_* is the traction force applied on one of *N_i_* pillars. Considering Eqs.(7), the intercellular force results in

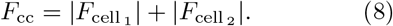

### Statistics

P-values between two groups were calculated using the two-sided Wilcoxon rank-sum test in MATLAB. Data sets were significantly different with probabilities of *p* ≤ 0.05 (*); p < 0.0001 (****); p > 0.05 (ns).

## Acknowledgments

We acknowledge Erik Danen (Leiden University) for providing the MV3 cell line. The authors thank the Austrian Science Fund (I 4971-N) for partially funding this work.

## Authors’ contributions

J.E. conducted and coordinated the research, performed analytic work, experiments, and data analysis. S.K., V.M. and S.P. manufactured the wafer for the CC-SD, J.E. and T.S. wrote the manuscript. T.S. supervised the project.

## Supplementary information

**Table S1.**
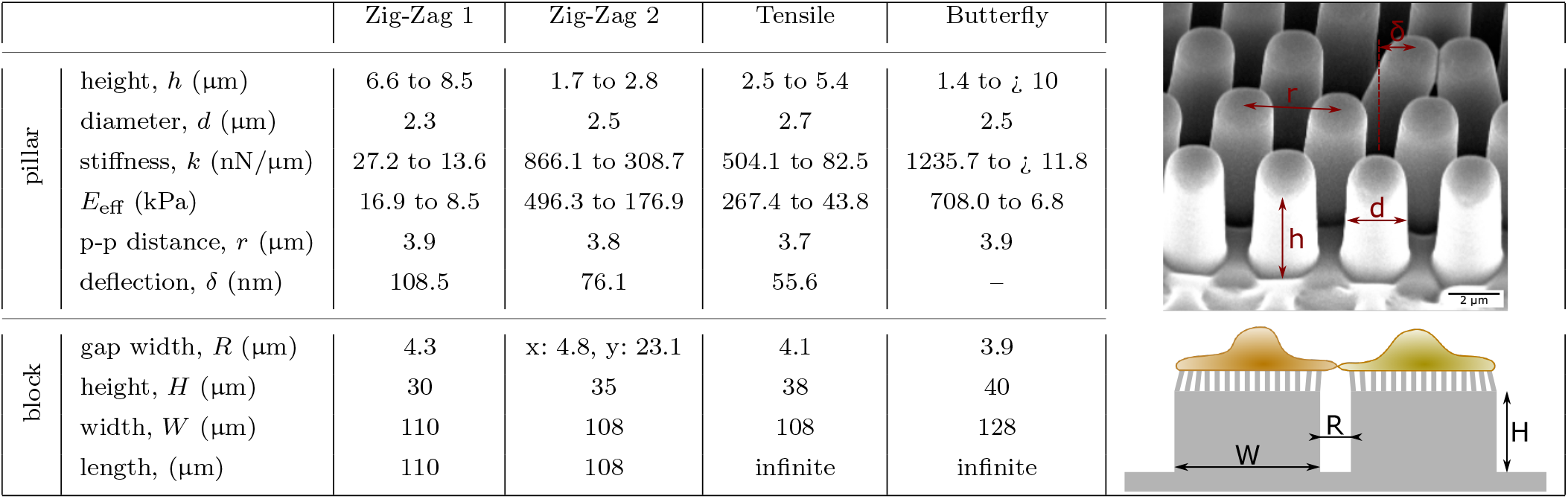
Cell-Cell Separation Device (CC-SD) specifications. This table summarizes all characteristics of the four different CC-SDs. The bending stiffness, *k*, was calculated with (1b) and the effective Young’s modulus, *E*_eff_ with (3). All other values were obtained by analyzing SEM images using ImageJ and Matlab software. The substrates are made of PDMS with a Young’s modulus of E = 2500 kPa.

**Fig. S1.**
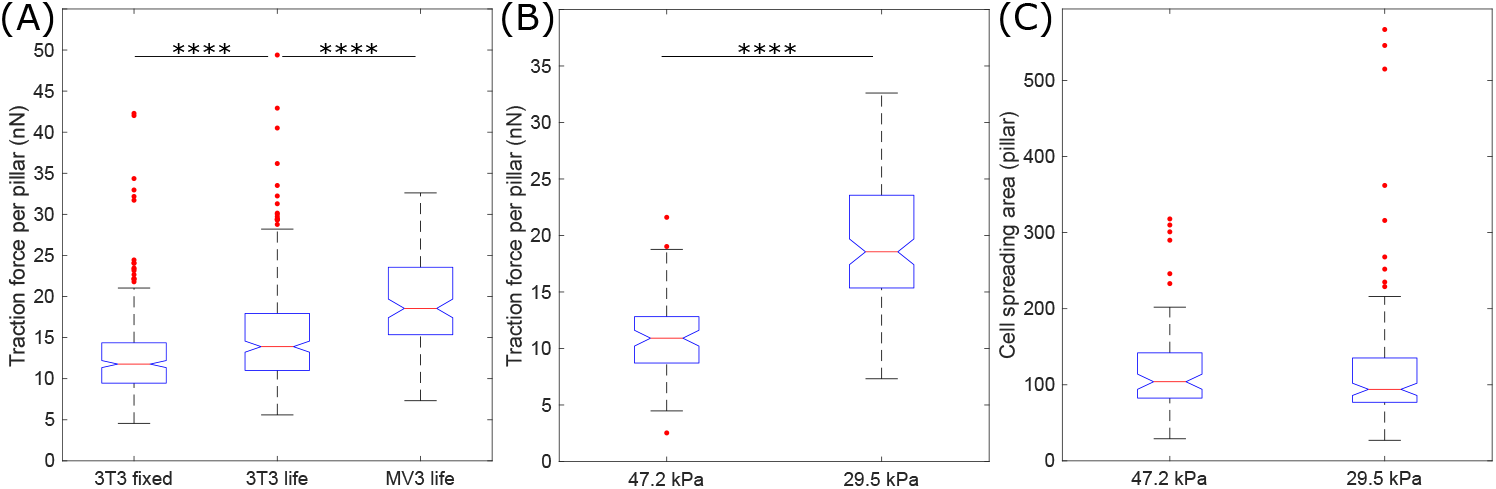
(**A**), Traction force analysis of single 3T3 fibroblasts (fixed and life) and MV3 cells cultured on micropillar arrays with Young’s modulus of 29.5 kPa (n_3T3-fixed_=321, n_3T3-life_=259). (**A,B**), Single MV3 cells cultured on elastic micropillar arrays with Young’s modulus of 47.2 kPa and 29.5 kPa (*n*_stiff_=85, *n*_soft_ = 128). (**A**), Cells apply larger traction forces on soft pillars compared to stiff. (**B**), The cell spreading areas are equal. Two-sided Wilcoxon rank sum test: **** p < 0.0001.

**Fig. S2.**
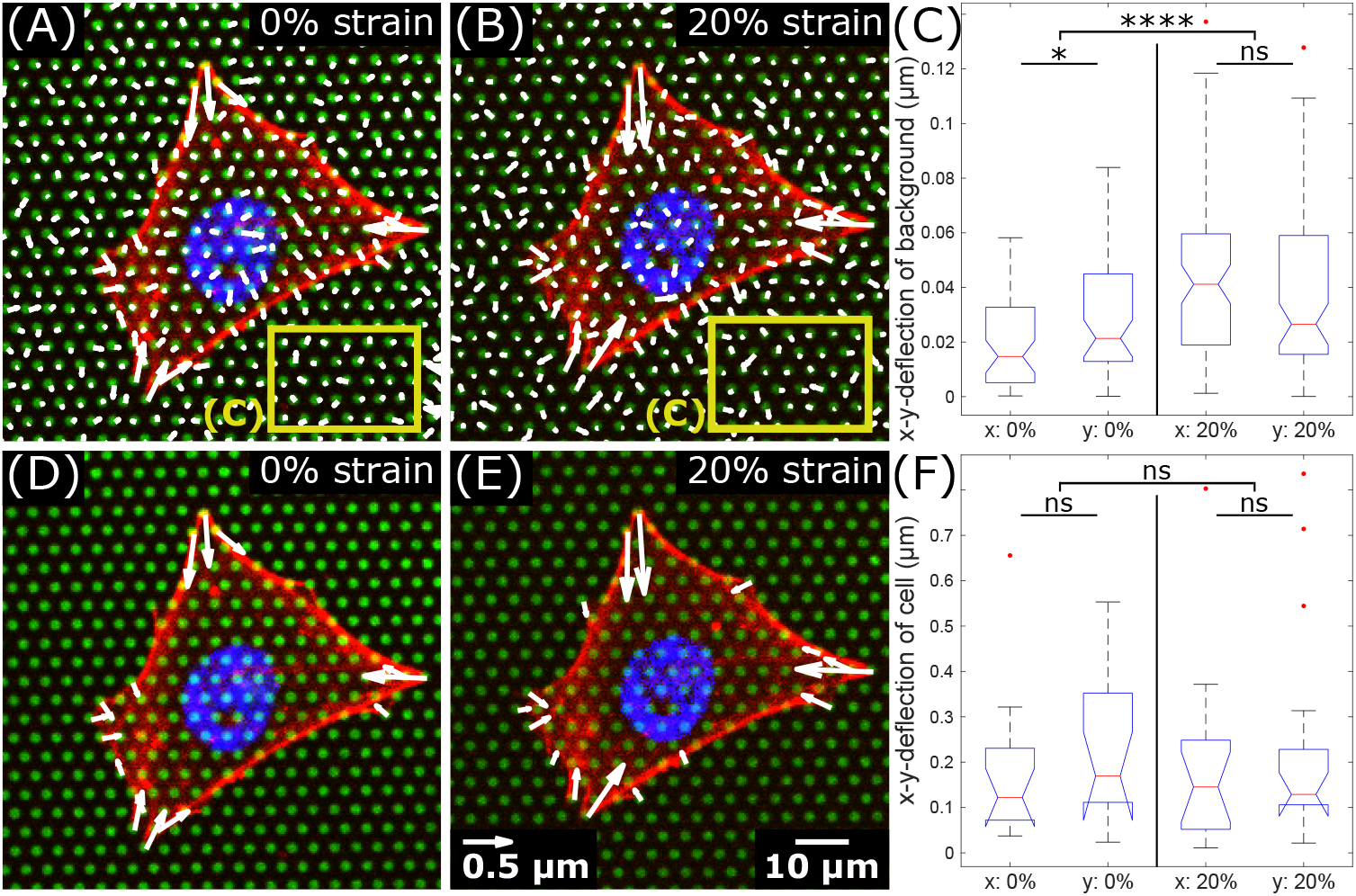
A single MV3 cell cultured on an elastic micropillar array with Young’s modulus of 47.2 kPa. (**A,B**), The deflection field of all pillars at 0% and 20% nominal substrate strain. The yellow frame represents the pillars considered for background analysis in (**C**). (**C**), Absolute deflections in x-and y-direction of pillars in a manual selected background region, i.e. excluding pillars covered by cells. In total, 54 and 82 pillars at 0% and 20% nominal strain, respectively, were considered. (**D,E**), Deflection of pillars caused by cell traction forces at 0% and 20% nominal substrate strain. Only deflections towards the center of mass of the cell are shown. Background forces were excluded. (**F**), Absolute deflections in x-and y-direction of pillars deflected by the cell. In total, 15 and 16 pillars at 0% and 20% nominal strain, respectively, were considered. Green: fibronectin-coated micropillars, red: F-actin, blue: nuclei. Two-sided Wilcoxon rank sum test: * p ≤ 0.05, **** p < 0.0001, ^*ns*^ p > 0.05.

**Fig. S3.**
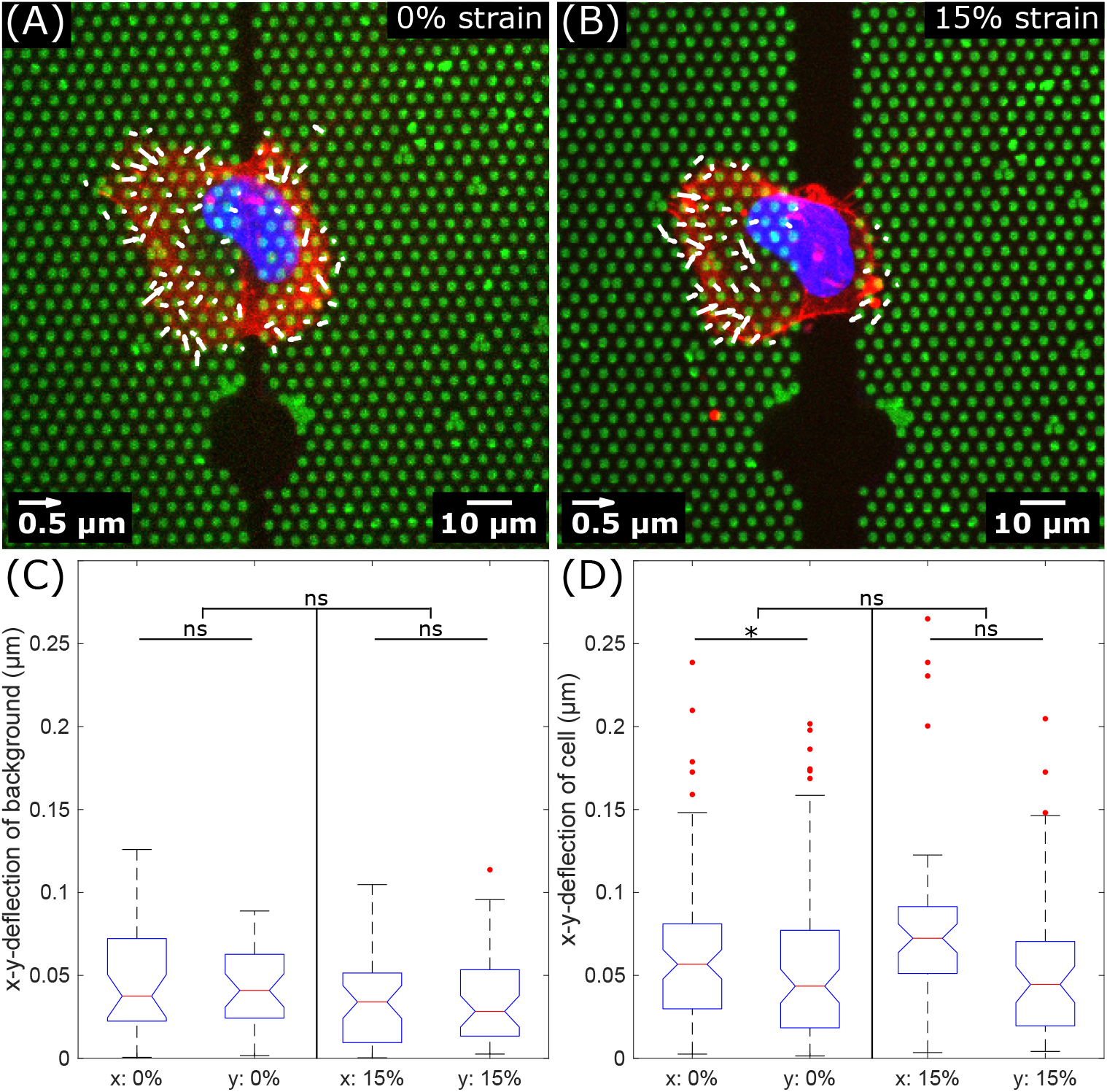
A single MV3 cell, cultured on the CC-SD, spread across the gap. (**A,B**), Deflection of pillars caused by cell traction forces at 0% and 15% nominal substrate strain. Only deflections towards the center of mass of the cell are shown. (**C**), Absolute deflections in x- and y-direction of pillars in a manual selected background region, i.e. excluding pillars covered by cells. At 0% strain, the mean x- and y-deflection was calculated to be 0.050 ± 0.036 μm and 0.043 ± 0.024 μm (n = 36 pillars), respectively, and at 15% strain, 0.034 ± 0.027 μm and 0.036 ± 0.028 μm (n = 44 pillars). (**D**), Absolute deflections in x-and y-direction of pillars deflected by the cell. At 0% strain, the mean x- and y-deflection below the cell was calculated to be 0.064 ± 0.046 μm and 0.06 ± 0.05 μm (n = 82 pillars), respectively, and at 15% strain, 0.08 ± 0.05 μm and 0.055 ± 0.045 μm (n = 53 pillars). Green: fibronectin-coated micropillars, red: F-actin, blue: nucleus. Two-sided Wilcoxon rank sum test: * p ≤ 0.05, ^*ns*^ p > 0.05.

**Fig. S4.**
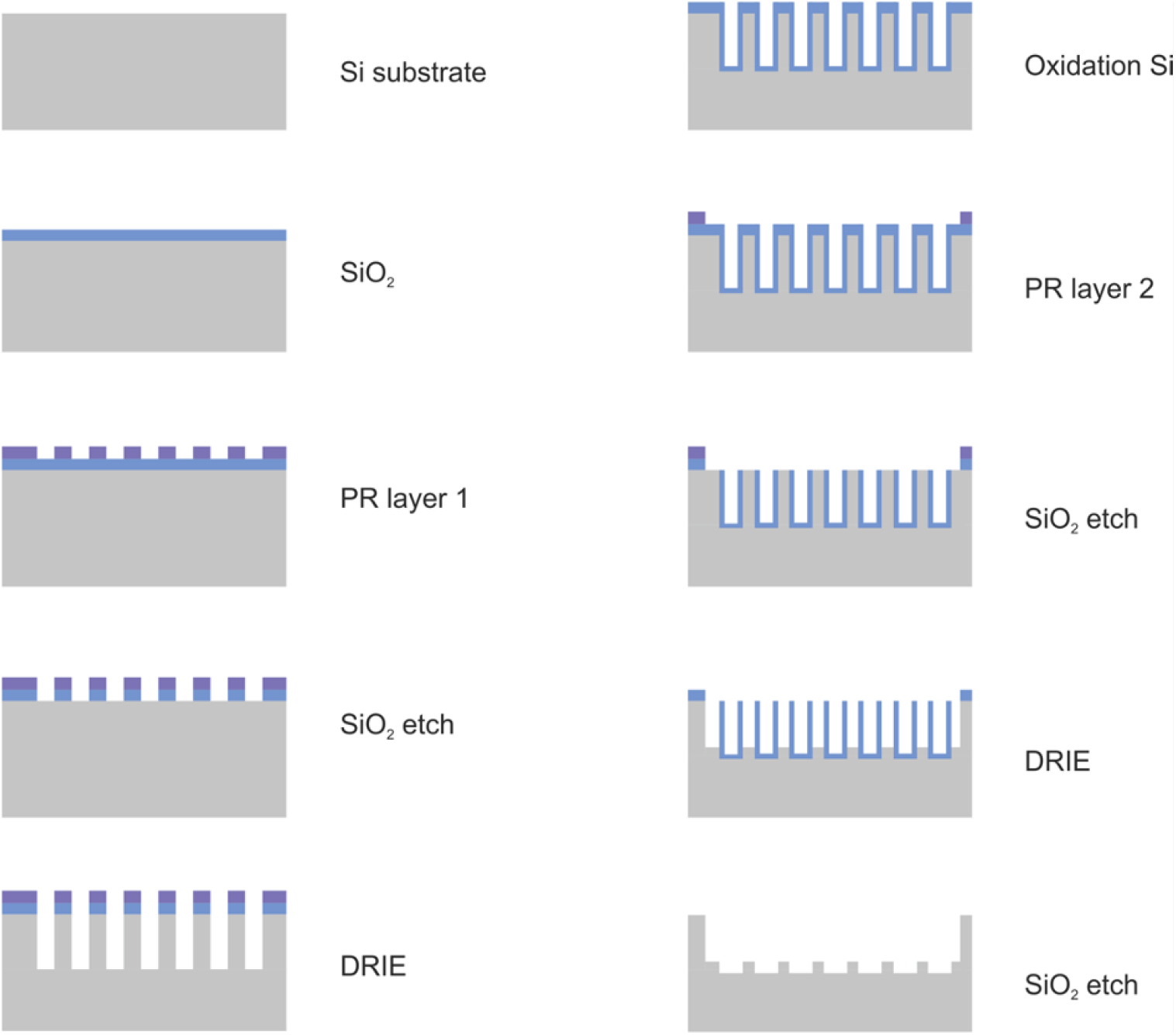
A silicon-wafer is oxidized by thermal oxidation (SiO_2_ in blue). Afterwards, the first photolithography step takes place, in which a thin layer of photoresist (PR in purple) is utilized to transfer the desire pattern and open the SiO_2_ mask. In this step, the holes that will be the PDMS pillars later are defined. After opening the SiO_2_ hard mask, the holes are transferred into the Si-substrate using the Bosch Process to etch deep into the substrate (DRIE). The remaining photoresist is removed and another SiO_2_ hard mask (blue) created by thermal oxidation. Next, the second photoresist layer (purple) is applied by spraying. This layer is used to separate the hole arrays (block height H) followed by opening the oxide mask by deep etching using the Bosch Process. In the final step, the remaining silicon oxide is removed by wet chemical etching (BOE).

